# Hypersensitivity controlled by mir-9a modulates female receptivity of *Drosophila melanogaster*

**DOI:** 10.1101/2024.07.29.605583

**Authors:** Xiaoli Zhang, Joshua A. Bagley, Yongwen Huang, Woo Jae Kim

## Abstract

Female *Drosophila melanogaster* undergo a complex behavioral transformation following mating, characterized by increased sensory sensitivity and altered reproductive behaviors. In this study, we investigated the role of miR-9a, a conserved microRNA, in regulating these post-mating changes. We found that miR-9a mutant females exhibited a hypersensitivity phenotype, with increased rejection of courting males, delayed onset of sexual receptivity, and abnormal mating termination behavior. This phenotype was associated with aberrant overgrowth of adult body wall sensory neurons, suggesting a link between neuronal hypersensitivity and reproductive behavior. To further elucidate the underlying mechanisms, we performed genetic interaction studies with *sens* and *bru2*, genes known to interact with miR-9a. We found that removing one copy of *sens* or *bru2* in miR-9a mutant backgrounds rescued the female rejection phenotype and normalized neuronal morphology. This suggests that miR-9a regulates sensory neuron development and female receptivity by modulating the expression of target genes like *sens* and *bru2*. Our findings reveal a novel role for miR-9a in regulating sensory hypersensitivity and reproductive behaviors in *Drosophila*. This research provides valuable insights into the molecular mechanisms underlying post-mating behavioral adaptations and neural development. Additionally, our findings highlight the potential of *Drosophila* as a model organism for investigating the role of miR-9 family members in neuronal specification and function, with implications for understanding sensory processing and neural plasticity in other organisms.

## INTRODUCTION

Hypersensitivity in sensory neurons refers to an increased sensitivity to stimuli, which can result in exaggerated or inappropriate responses to normal sensory input (Isaacs and Riordan 2020).

This can manifest as pain, itching, or discomfort in response to stimuli that would not normally be perceived as painful or unpleasant (Latremoliere and Woolf 2009; Gangadharan and Kuner 2013). Hypersensitivity can occur in various forms and is often associated with certain neurological disorders, injuries, or inflammatory conditions (Ren and Dubner 2008; Pinho-Ribeiro et al. 2017). Research into the causes and mechanisms of hypersensitivity in sensory neurons is important for developing new treatments to alleviate chronic pain and improve the quality of life for those affected by these conditions.

Hypersensitivity can be a result of changes in the nervous system known as neuroplasticity (Petersen-Felix and Curatolo 2002; Latremoliere and Woolf 2009; Gangadharan and Kuner 2013). This can involve altered gene expression in sensory neurons, changes in the morphology and function of these neurons, and modifications in the way they interact with other cells in the nervous system (Woolf and Salter 2000). In some cases, hypersensitivity can become chronic due to central sensitization, a process where the central nervous system becomes hyper-excitable and develops an increased sensitivity to incoming sensory signals. This can result in long-lasting pain even after the initial injury or inflammation has resolved (Salter 2010).

After mating, female *Drosophila melanogaster* undergo a marked transformation in behavioral response, exhibiting an enhanced sensitivity to a broad array of sensory inputs. This altered sensory perception is a component of a complex suite of post-mating changes that optimize the female’s reproductive strategy (Yang et al. 2009; Zhu et al. 2014; Hussain et al. 2016; Bath et al. 2017). Enhanced sensory reactivity represents a pivotal component of the post-mating behavioral syndrome in female *Drosophila melanogaster*. This adaptive sensitization is hypothesized to confer selective advantages by facilitating the evasion of predators and/or by aiding in the detection of optimal oviposition sites, thereby enhancing the female’s reproductive success (Hollis et al. 2019).

MicroRNAs (miRNAs) are a class of endogenous small non-coding RNAs that predominantly function in the post-transcriptional regulation of gene expression. These molecules exert their regulatory effects by annealing to the complementary sequences within the 3’ untranslated region (3’ UTR) of target messenger RNAs (mRNAs), thereby facilitating either the degradation of the mRNA transcript or the suppression of protein synthesis, depending on the degree of complementarity and other context-dependent factors (Bartel 2009; Brodersen and Voinnet 2009). miRNAs are implicated in diverse brain functions including development, cognition, and synaptic plasticity (Smalheiser and Lugli 2009; Cohen et al. 2011; Aksoy-Aksel et al. 2014; Ye et al. 2016; Mohammadi et al. 2022).

Although the role of miRNAs in neural development and plasticity is well-documented, their specific contributions to the modulation of sensory hypersensitivity following mating in female insects have not been extensively explored. Studies using genetic tools to manipulate miRNA expression in specific neurons or at specific times after mating could help to elucidate the precise mechanisms by which miRNAs contribute to these complex behavioral and physiological changes. Such studies are poised to elucidate the molecular underpinnings of post-mating sensory adaptations and their implications for reproductive success.

miR-9a and its mammalian ortholog, miR-9, are pivotal regulators of post-transcriptional gene expression, with diverse roles in development, cellular differentiation, and disease progression (Li et al. 2006; Biryukova et al. 2009; Cassidy et al. 2013; Li et al. 2013; Yatsenko and Shcherbata 2014; Cassidy et al. 2015; Suh et al. 2015; Daniel et al. 2017; Katti et al. 2017; Gallicchio et al. 2020; Subramanian et al. 2021). Although miR-9a is multifunctional, miR-9a was initially recognized for its essential function in neural development, particularly in the precise specification of sensory organ precursors (SOPs) (Li et al. 2006; Parrish et al. 2006).

Notably, miR-9a exhibits a strong genetic interaction with the senseless (sens) gene in controlling SOPs formation (Li et al. 2006). The miR-9a predominantly influences the development of multidendritic sensory neurons and is crucial for the correct morphogenesis of their dendritic arbors (Parrish et al. 2006; Wang et al. 2016).

In this study, we demonstrate that miR-9a regulates female receptivity in *Drosophila melanogaster* through a sens-dependent pathway. Non-mated females with mutated miR-9a exhibit a hypersensitivity phenotype, characterized by increased rejections of males, akin to the behavior of mated females. This behavioral shift is correlated with the overgrowth of adult body wall sensory neurons. Our findings delineate a novel role for miR-9a in modulating neuronal hypersensitivity and provide insights into the molecular mechanisms governing post-mating behavioral changes.

## RESULTS

### Female *Drosophila melanogaster* with mutations in *miR-9a* exhibit rejection of courting males

It is established that internal sensory neurons expressing the *pickpocket* (*ppk*) gene mediate the post-mating behavioral switch through the binding of sex peptide to its receptor (Häsemeyer et al. 2009; Yang et al. 2009). Given that miR-9a is implicated in the development of sensory organ precursors (SOPs) (Li et al. 2006; Wang et al. 2016), including ppk-positive neurons, we propose that miR-9a may regulate female receptivity by modulating the development of ppk-expressing neurons in females.

Both *miR-9a* mutant strains, *mir-9a^J22^* and *mir-9a^E39^*, displayed a significant decrement in receptivity among virgin females across both 20-minute and 120-minute observation periods (Fig. 1A-B and Fig. S1A-B). Temporal analysis of receptivity scores definitively revealed that miR-9a mutant females exhibit a substantial delay in the onset of sexual receptivity (Fig. 1C and Fig. S1C). Relative to wild-type control females, miR-9a mutant females exhibited a significantly higher rejection rate within a 5-minute observation window (Fig. 1D). Moreover, miR-9a mutant females that initially accepted a male often subsequently terminated the mating attempt (Fig. 1E). This behavior, wherein a female accepts a male and then rapidly rejects accepted male within seconds, is a phenotype that is infrequently observed in wild-type females (Movies 1-4).

**Figure 1.**
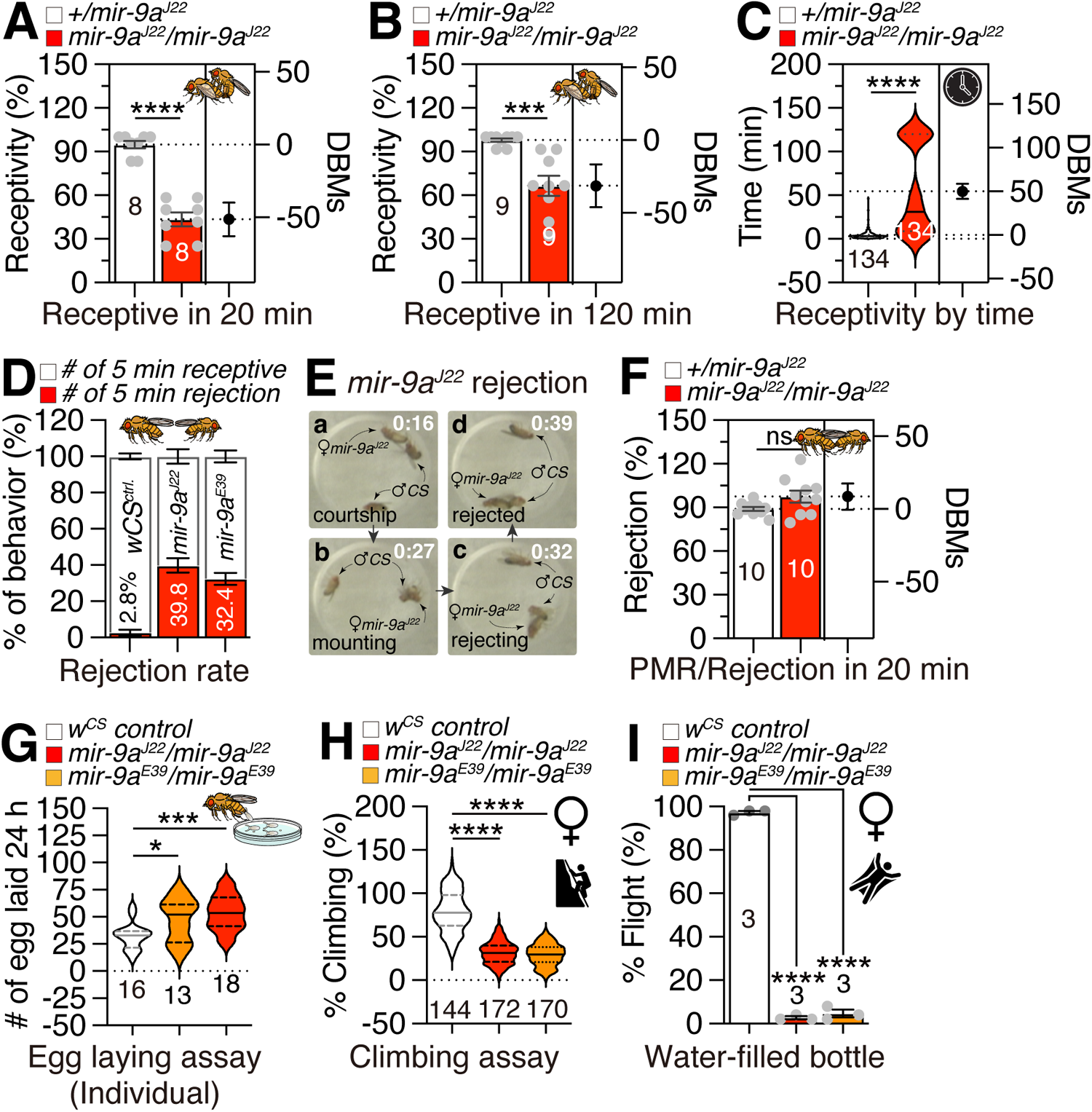
*miR-9a* mutations lead to a specific increase in rejection behavior among virgin females. (A-B) Receptivity of virgin females, with the score for receptivity behavior as well as genotype of experimental animals given above the graph. In each assay, one female of the indicated genotype was confronted with two naive males in a small chamber. Females were scored as receptive when they mated within 20 minutes (A) and 120 minutes (B). Numbers in parentheses are the numbers of females tested. DBMs represent difference between means. The mean value and standard error are labeled within the dot plot (black lines). Asterisks represent significant differences, as revealed by the Student’s t test (* p<0.05, ** p<0.01, *** p<0.001). The same notations for statistical significance are used in other figures. (C) The temporal pattern of virgin female receptivity to males courtship across the duration of the observation period. (D) The Receptivity assays of *wCS* females, *mir-9a^J22^/mir-9a^J22^* homozygous mutant and *mir-9a^E39^/mir-9a^E39^* homozygous mutant in 5 minutes. (E) Females harboring a *mir-9a^J22^* mutation initially accepted male courtship but subsequently terminated the mating attempt. (F) Rejection of mated females, with the score for rejection behavior as well as genotype of experimental animals given above the graph. PMR means post-mating reaction. (G) Egg laying. For egg laying, 10 females of the appropriate genotype were aged in vials for 4–5 days. Then three or five females were transferred to a vial with grape media and allowed to lay eggs for 24 hr at 25°C. The number of eggs was divided by the number of flies in the vial to give a measure of egg laying. (H) Climbing assays of females. 40–50 flies were placed in an empty vial and were tapped to the bottom of the tube. After tapping of flies, we recorded 10 s of video clip. This experiment was done five times at 5-min intervals. With recorded video files, we captured the position of flies 10 s after tapping the vial. (I) Flight assays of females. For each assay, fifty flies were gently introduced into a water-filled jar. The jar was then tapped to stimulate the flies, and the number of flies that escaped from the water was counted. The escape ratio was calculated to determine the effectiveness of the flies’ flight response.

Post-mating rejection of male courtship is a conserved element of female post-mating responses (PMR) in *Drosophila melanogaster* (Chapman et al. 2003; Kubli 2003; Yapici et al. 2008).

While miR-9a mutant virgin females displayed a significantly elevated rejection rate compared to controls, the rejection rate of mated miR-9a mutants was indistinguishable from that of control mated females (Fig. 1F and Fig. S1D), indicating that the PMR in miR-9a mutants is intact.

Notably, mated miR-9a mutants deposited a greater number of eggs than controls (Fig. 1G and Fig. S1E), suggesting that miR-9a specifically modulates egg-laying behavior rather than rejection behavior within the PMR. Furthermore, miR-9a mutants exhibited a marked reduction in climbing (Fig. 1H) and flight (Fig. 1I; Movies 5-6) abilities, indicative of impaired muscle contraction, consistent with previous reports (Katti et al. 2017). Collectively, these findings indicate that miR-9a mutations lead to a specific increase in rejection behavior among virgin females, while the rejection responses of mated females remain unaffected.

### Male fruit flies with mutations in miR-9a display abberent courtship behaviors, and the larvae of these mutants exhibit locomotor defects

To investigate the role of miR-9a in male sexual behavior, we assessed the courtship index and observed a significant reduction in courtship activity among miR-9a mutant males compared to controls (Fig. 2A). Notably, a substantial proportion of miR-9a mutant males displayed a unique courtship behavior characterized by bilateral wing vibration (Fig. 2B), in contrast to the typical unilateral wing vibration observed in controls (Fig. 2C). Approximately two-thirds of miR-9a mutant males exhibited this double wing vibration phenotype (Fig. 2D and Movies 7-8), indicating that miR-9a mutation leads to specific courtship defects in male flies. Similar to the locomotor deficits observed in mutant females, miR-9a mutant males also exhibited reduced climbing and flight abilities (Fig. S2A-B), suggesting a generalized impact of miR-9a mutation on locomotor behavior.

**Figure 2.**
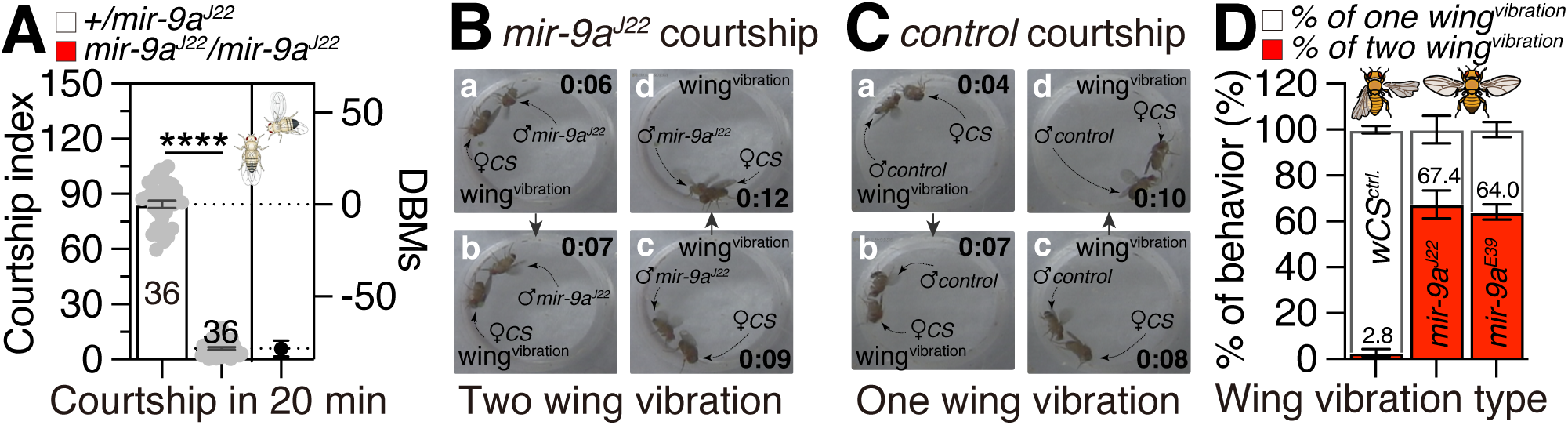
Males with *mir-9a* mutation display abberent courtship behaviors. (A) Courtship assays of males. Once courtship began, courtship index was calculated as the fraction of time a male spent in any courtship-related activity during a 10 min period or until mating occurred. Mating initiation is the time after male flies successfully mounted on female. (B) *miR-9a^J22^* males displayed a unique courtship behavior characterized by bilateral wing vibration. (C) *CS* males displayed a typical courtship behavior characterized by unilateral wing vibration. (D) Wing vibration types of males. White box represents the percentage of unilateral wing vibration and red box represents the percentage of bilateral wing vibration.

In line with these adult locomotion data, third instar larvae of miR-9a mutants showed impaired locomotion (Fig. S2C-E) and exhibited an increased frequency of turning behaviors compared to controls (Fig. S2F and Movies 9-10), indicating that the miR-9a mutation elicits a unique behavioral profile, distinct from a generalized impairment of health. These findings collectively suggest that miR-9a mutation induces a range of sensory-motor-related phenotypes, from larvae to adults, across both sexes.

### In miR-9a mutant flies, the adult body wall neurons undergo aberrant overgrowth

The conservation of miR-9 across evolutionary scales, from flies to humans, at the nucleotide level underscores its functional significance, despite variations in expression patterns. Studies across various model organisms have revealed that miR-9 can influence neurogenesis through its regulatory role in both neural and non-neural cell lineages (Yuva-Aydemir et al. 2011). In *Drosophila*, miR-9a is essential for the accurate specification of neural progenitor cells in non-neural lineages. For instance, the loss of miR-9a activity leads to the formation of ectopic sensory neurons in embryos, larvae, and adults (Li et al. 2006).

To assess whether the female mating rejection phenotype is associated with aberrant sensory neuron development, we analyzed the female abdominal body wall neurons, which derive from the larval sensory neurons that are regulated by miR-9a (Li et al. 2006). Consistent with previous findings, *ppk*-positive larval body wall sensory neurons in miR-9a mutants exhibit ectopic sensory neurons with exceptionally branched dendrites (Fig. S3A-B). Remarkably, adult body wall neurons also display ectopic sensory neurons with overgrown dendrites in both the ventral and dorsal abdomen (Fig. 3A-B and Fig. S3C-D). Quantification of neurite morphology revealed a significant increase in the number of branches and junctions in miR-9a mutant body wall neurons (Fig. 3C-D and Fig. S3E-F), although the average length of the branches is comparable between the mutant and control groups (Fig. 3E and Fig. S3G). The collective data indicate that miR-9a is instrumental in the regulation of adult body wall neuronal growth.

**Figure 3.**
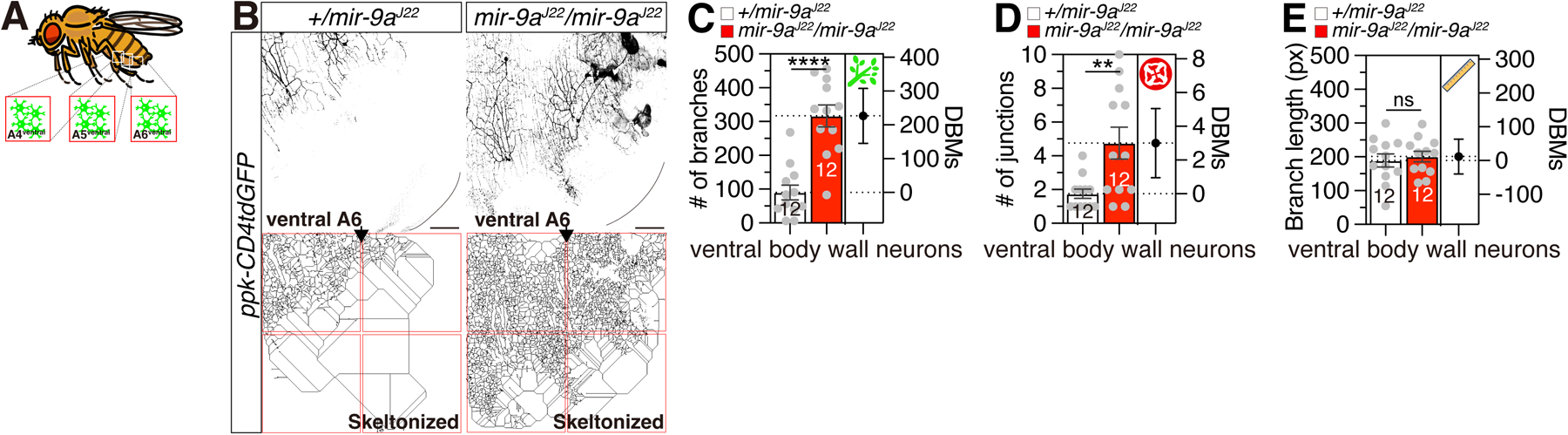
The adult ventral body wall neurons undergo aberrant overgrowth in *mir-9a* mutant flies. (A) Location of *Drosophila* adult ventral neurons. (B) Ventral A6 neurons expressing *ppk-GAL4* together with *UAS-mCD4GFP* in *mir-9a^J22^/ +* and *mir-9a^J22^/mir-9a^J22^* in adult female. The bottom pictures are skeletonized from the top pictures. (C-E) Quantification of neurite morphology for *miR-9a* mutant body wall neurons in branches (C), junctions (D) and branch length (E).

### *Sensless* (*sens*) and *bruno 2* (*bru2*) mutations rescue the phenotypic defects of miR-9a mutant flies

It has been reported miR-9a mutants exhibit sensory bristle defects on the notum, and miR-9a interacts genetically with the *sens* gene (Li et al. 2006). To determine if adult body wall neurons are also affected by miR-9a mutation, we conducted genetic interaction experiments. In a *miR- 9a^J22^* homozygous mutant background, the removal of one copy of *sens* (*sens^E58^/+*) resulted in a significant rescue of the receptivity defects, matching those of control females (Fig. 4A and Fig. S4A). This genetic rescue of the receptivity phenotype is correlated with the normalization of the adult body wall neuronal phenotype, which displays normal sensory neurons with typical neurite growth (Fig. 4B-E).

**Figure 4.**
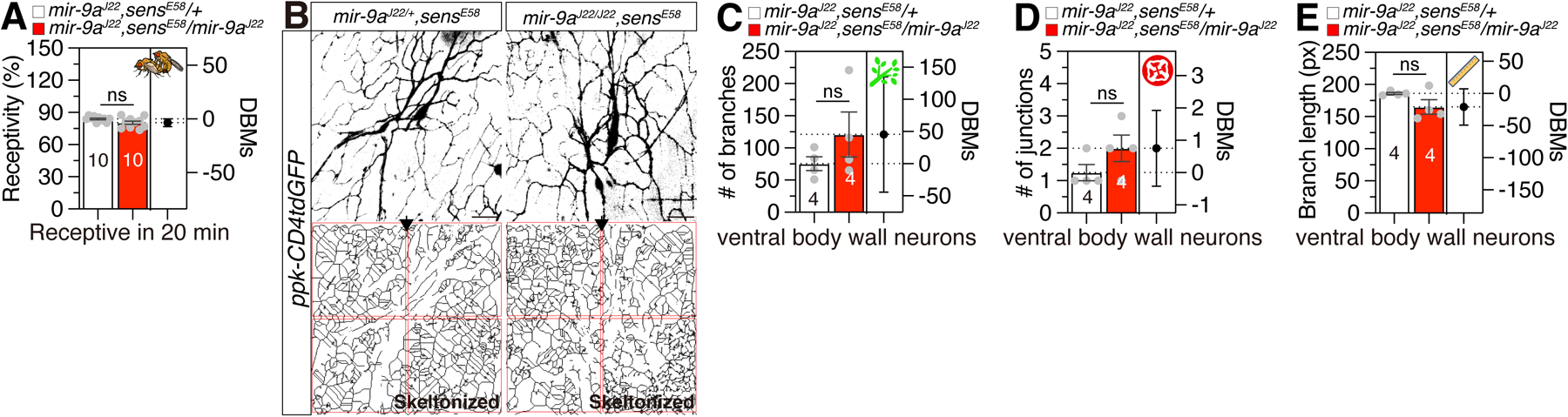
*sens* and *bru2* mutations rescue the phenotypic defects of *miR-9a* mutant flies. (A) Receptivity of virgin females, with the score for receptivity behavior as well as genotype of experimental animals given above the graph. (B) Ventral neurons expressing *ppk-GAL4* together with *UAS-mCD4GFP* in *mir-9a^J22^/ +, sens^E58^* and *mir-9a^J22^/mir-9a^J22^, E58* female. The bottom pictures are skeletonized from the top pictures. (C-E) Quantification of neurite morphology for venral body wall neurons in branches (C), junctions (D) and branch length (E). Genotype of experimental animals given above the graph.

Through bioinformatic analysis, we identified several miR-9a target genes that contain miR-9a target sequences in their untranslated regions (UTRs) of mRNA. We conducted similar genetic interaction tests as with *sens* mutant and found that the removal of one copy of *bru2* (*bru2/+*) completely rescued the miR-9a mutant receptivity phenotype (Fig. 5A). Overexpression of bru2 in sensory neurons did not alter the receptivity phenotype (Fig. 5B), indicating that *bru-2* is not a direct target of miR-9a in sensory neurons. The *Drosophila* gene *bru2* is predicted to facilitate mRNA 3’-UTR binding activity and is involved in the negative regulation of translation (Öztürk-Çolak et al. 2024). These findings collectively suggest that miR-9a guides sensory neuron specification in the adult body wall and controls virgin female receptivity by regulating the expression of various miR-9a target genes, including *sens* and *bru2*.

**Figure 5.**
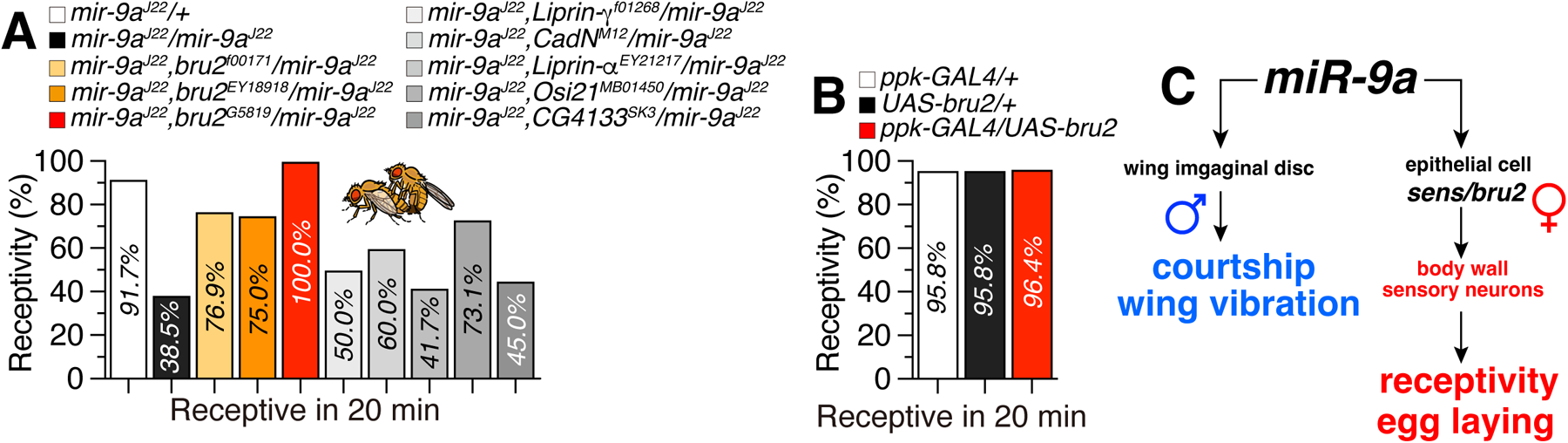
*miR-9a* guides sensory neuron development in the adult body wall and modulates virgin female receptivity through target gene regulation. (A-B) Receptivity of virgin females in 20 minutes, with the score for receptivity behavior as well as genotype of experimental animals given above the graph. (C) Schematic diagram of mir-9a involved in sexual behavior in females and males.

## DISCUSSION

Our study reveals the critical role of miR-9a in regulating reproductive behaviors and neural development in *Drosophila melanogaster*. Female miR-9a mutants display a pronounced phenotype characterized by increased rejection of courting males, delayed onset of sexual receptivity, and abnormal mating termination behavior. While the post-mating rejection behavior of mated females remains intact, they lay more eggs, suggesting specific effects on egg-laying (Fig. 1). Similar locomotor deficits are observed in both sexes, highlighting the generalized impact of miR-9a mutation on motor function. In male flies, miR-9a mutation leads to reduced courtship activity and the emergence of an abnormal bilateral wing vibration phenotype (Fig. 2). Further investigation of sensory neuron development reveals ectopic overgrowth of sensory neurons in both larval and adult stages, indicating a disruption in neuronal specification and growth (Fig. 3). Genetic interaction experiments with *sens* and *bru2*, genes involved in sensory bristle development and mRNA translation regulation, respectively, demonstrate that removing one copy of these genes in miR-9a mutant backgrounds rescues the receptivity defects and normalizes neuronal morphology (Fig. 4 and 5). These findings collectively suggest that miR-9a plays a crucial role in regulating the development of sensory neurons and female receptivity by modulating the expression of target genes such as *sens* and *bru2*. This study provides valuable insights into the molecular mechanisms underlying reproductive behaviors and neural development in *Drosophila*, and potentially in other organisms as well.

In our investigation, we have elucidated a heretofore unreported target gene of miR-9a, designated *bru2*, which, upon deletion of a single allele, can rescue the female rejection phenotype induced by miR-9a mutation. The *Drosophila melanogaster bru2* gene is inferred to possess mRNA 3’-UTR binding activity and is proposed to participate in mRNA splice site recognition, the negative regulation of translation, and the regulation of alternative mRNA splicing, mediated through the spliceosome. Despite these predictions, the precise molecular function of bru2 remains to be fully elucidated. The human orthologs of bru2, CELF1 (CUGBP Elav-like family member 1) and CELF2 (CUGBP Elav-like family member 2), have been implicated in developmental and epileptic encephalopathy 97 (Itai et al. 2021). Notably, a constellation of repressed expression of let-7g, miR-9, and miR-135a, alongside elevated expression of the RNA splicing factor CELF1, has been correlated with a murine heart model of myotonic dystrophy (MD) (Misra et al. 2020). These findings suggest that the newly characterized genetic interplay between miR-9a and bru2 may serve as a prototypical model for investigating the role of human miR-9 in neuronal specification.

Our research revealed that mutation of miR-9a results in the overgrowth of sensory neurons in adult *D. melanogaster* females, which in turn elicits rejection of courting males by virgins. The miR-9a mutant females exhibited a hypersensitivity phenotype attributable to this neuronal overgrowth. The study of hypersensitivity in *D. melanogaster* is a field of inquiry that holds profound implications for comprehending the foundational mechanisms underpinning sensory perception and neural plasticity. In *D. melanogaster*, post-mating modifications in female behavior encompass enhanced sensory reactivity, a phenomenon considered adaptive as it facilitates the evasion of predators and the selection of appropriate oviposition sites (Chapman et al. 2003; Ram and Wolfner 2007; Yapici et al. 2008; Gligorov et al. 2013; Corbel et al. 2022).

This augmented sensitivity is correlated with the overgrowth of adult body wall sensory neurons, and alterations in miR-9a expression via genetic manipulations can modulate these responses.

The analysis of hypersensitivity in this fly model provides critical insights into the molecular and neural substrates that regulate sensory processing and the evolution of complex behavioral outputs.

Notably, female *ppk*-positive body wall neurons have been identified as key neurons underlying the heat hypersensitivity phenotype (Gu et al. 2022), indicating that adult body wall neurons are evolutionarily conserved in mediating adaptive hypersensitivity. Our discovery establishes a novel *Drosophila* genetic model for the investigation of miR-9 family-mediated pathogenesis across various neuronal contexts.

## METHODS

### Quantification of Neurite Branching

To quantify neurite branching, we utilized the ’skeleton’ function in ImageJ, following a detailed step-by-step process to ensure accurate measurement of branch numbers and lengths. All specimens were imaged under identical conditions. First, images of neurons were imported into ImageJ by launching the software and dragging the image into the workspace. The images were then converted to an 8-bit format by selecting the image, navigating to the “Image” menu, choosing “Type,” and selecting “8-bit.” Next, the images underwent threshold adjustment to separate the neuron branches from the background. This involved selecting “Image” from the menu bar, choosing “Adjust,” and then “Threshold.” The “Dark Background” option was checked, the color was changed to red, and the threshold slider was adjusted until all branches were highlighted in red. After threshold adjustment and branch connection, the images were converted to a binary format. This was done by selecting “Process” from the menu bar, choosing “Binary,” and then “Make Binary,” ensuring that “Threshold pixel to foreground color” and “Remaining pixel to background color” options were checked with a “Black foreground, white background” setting. The binary images were then skeletonized to produce a simplified representation of the neurite branches. This involved selecting “Process” from the menu bar, choosing “Binary,” and then “Skeletonize.” The resulting skeleton images were analyzed using the Analyze Skeleton plugin. This process included selecting “Analyze” from the menu bar, choosing “Skeleton,” and then “Analyze Skeleton.” The settings for the analysis included keeping the “Prune cycle method” set to “None,” unchecking the “Prune ends” and “Exclude ROI from pruning” options, and checking the options to show the longest shortest path, branch labels, junctions, and endpoints. All specimens were imaged under identical conditions.

### Predicting of miR-9a Targets in *Drosophila* Genome

We used miRBase provided targetscan analysis to identify miR-9a potential targets (Kozomara et al. 2019). The result can be checked with this stable link. https://www.targetscan.org/cgi-bin/targetscan/fly_12/targetscan.cgi?mirg=dme-miR-9a

### Adult Flight Assay

Flies were subjected to a ’flight assay’ to evaluate their escape response from water. 50 flies were gently introduced into a water-filled jar. The jar was then tapped to stimulate the flies, and the number of flies that escaped from the water was counted. The escape ratio was calculated to determine the effectiveness of the flies’ flight response. For visual reference, please check the video files labeled 5-6, which represent the phenotypes of control and mutant flies during the assay. Our experiment was conducted at 25°C under normal light conditions.

### Receptivity Assay and Egg Laying Assay

For receptivity assays, females and males were housed in small groups of 3–4 flies. In order to assess female receptivity, one female was put into a small (1 cm × 1 cm) chamber together with two naive *Canton S* males. The female was scored as receptive if it mated within 20 min. In the remating assays, a female was mated in a receptivity assay and then again examined in another receptivity assay 24 hours later. Assays were usually performed within the first three hours of the subjective day. All experimental procedures were conducted using 36-well plates, and each assay was replicated the specified number of times to ensure statistical robustness. The calculated percentages of receptivity and rejection were then subjected to statistical analysis via Student’s t-test for significance.

For egg laying assays, 10 females of the appropriate genotype were aged in vials for 4–5 days. Then three or five (as indicated in the figure legends) females were transferred to a vial with grape media and allowed to lay eggs for 24 hours at 25°C. The number of eggs was divided by the number of flies in the vial to give a measure of egg laying. For assays of egg laying by mated females, females of the respective genotype were mated with *Canton S* males for 2–3 hours on the day before the experiment, with 10 females and 20 males per vial. All data are given as average ± SEM, significance levels were calculated with the Student’s t test.

### Courtship Assay

The courtship assay was conducted according to established protocols as previously reported (Lee et al. 2023), under standard light conditions in circular courtship arenas with a diameter of 11 mm, between noon and 4 p.m. Courtship latency was defined as the interval from the introduction of the female to the initiation of the first overt male courtship behavior, such as orientation and wing extensions. Following the onset of courtship, the courtship index was determined as the proportion of time the male engaged in courtship-related activities over a 10- minute period or until copulation occurred.

### Video Recording of Larval Locomotor Behaviors

Video recordings of gross path morphology were made with a digital video (DV) camera (Canon GL1) in an environment room maintained at 25°C and 70% humidity. DV movies were captured with IMOVIE 2.0 on a 500-MHz Apple iMac and digitized with QUICKTIME 4.0 at 29.97 frames per second (fps). Low-magnification videos were recorded for 2 min or until the larva left the 50-cm^2^ field of the camera. High-magnification videos (15-cm^2^ field) were recorded with an Olympus OLY-200 camera mounted on an Olympus SZX9 microscope connected to a VCR (Samsung VR5599). Peristalsis was recorded until 10 peristaltic waves during linear locomotion were completed. Mutants frequently had discontinuous bouts of peristalsis, and repositioning of the plate was necessary, but this did not affect the motion analysis. VCR recordings were captured with adobe premier 5.0 on a Macintosh G4 at 15 fps.

### Climbing Assay

For climbing assay, we modified the conventional RING assay (Gargano et al. 2005) and reported in our previous reports (Miao et al. 2024; Zhang et al. 2024). In brief, 40-50 aged flies were placed in an empty vial and were tapped to the bottom of the tube. We used 5 days old adults. After tapping of flies, we recorded 10 seconds of video clip. This experiment was done five times with 5-minute intervals. With recorded video files, we captured the position of flies 10 seconds after tapping the vial. This captured image file was then loaded in ImageJ to perform particle analysis. For quantifying the location of flies inside a vial, we used the “analyze particles” function of ImageJ (Grishagin 2015). The position of pixels was normalized by height of vial then only the particles above the midline (4 cm) of vial were counted.

### Adult Body Wall Neuron Live Imaging

Adult body wall was dissected in PBS at room temperature. We imaged GFP fluorescence in living animals by mounting them in silicon oil (Shin-Etsu). Maximum projections of z-stacks were used in all cases. Images were adjusted for brightness and contrast with Adobe Photoshop (Adobe Systems, San Jose, CA) (Yasunaga et al. 2015). For visualization of dendrites, we labeled neurons with ppk-GAL4; UAS-mCD8GFP, ppk-CD4-tdGFP, or ppk-CD4-tdTom and imaged GFP and RFP fluorescence in living animals by mounting them in silicon oil (Shin-Etsu). Maximum projections of Z-stacks were used in all cases. The dendrite length was measured by using the ImageJ. For the quantitative analyses, we focused on the dendrites in segments A4, A5 and A6, since these neurons exhibit similar and consistent dendrite branch lengths and branch points. For quantification of the total branch length and the branch points, we used skeleton analysis as described above.

### Fly Stocks and Husbandry

*Drosophila melanogaster* were raised on cornmeal-yeast medium at similar densities to yield adults with similar body sizes. Flies were kept in 12 h light: 12 h dark cycles (LD) at 25°C (ZT 0 is the beginning of the light phase, ZT12 beginning of the dark phase) except for some experimental manipulation. Wild-type flies were *Canton-S* (*CS*) and *Cantonized w1118*.

Following lines used in this study, *Canton-S* (#64349), *ppk-CD4-tdGFP* (# 35842), *sens^E58^* (#5312), *bru2^f00171^* (#18300), *bru2^EY18918^* (#22296), *bru2^G5819^* (#27190), *Liprin-*γ *^f01268^* (#18421), *CadN^M1^*^2^ (#229), *Liprin-α ^EY21217^* (#22459), *Osi21^MB0145^ ^0^*(#23186), *ppk-GAL4* (#32078) were obtained from the Bloomington *Drosophila* Stock Center at Indiana University. *Osi21^MB01450^* (#M2L-3116) was obtained from National Institute of Genetics Fly Stocks. We thank Dr. Fen-Biao Gao (Gladstone Institute of Neurological Disease and Department of Neurology, University of California at San Francisco) for sharing *mir-9a^J22^* and *mir-9a^E39^ lines.* We thank Dr. Paul M. Macdonal (Department of Molecular Biosciences, Institute for Cellular and Molecular Biology, The University of Texas at Austin, Austin, Texas, United States of America) for sharing *UAS- bru2* line.

### Statistical Analysis

Statistical analysis of receptivity assays is similar with mating duration assay was described previously (Lee et al. 2023). More than 10 females for 1 group were used for receptivity assay. Statistical comparisons were made between groups that were control group and experimental group within each experiment. As receptivity assays of females showed normal distribution (Kolmogorov-Smirnov tests, p > 0.05), we used two-sided Student’s t tests. The mean ± standard error (s.e.m) (**** = p < 0.0001, *** = p < 0.001, ** = p < 0.01, * = p < 0.05). All analysis was done in GraphPad (Prism). Individual tests and significance are detailed in figure legends.

Besides traditional t-test for statistical analysis, we added estimation statistics for all receptivity assays and two group comparing graphs. In short, ‘estimation statistics’ is a simple framework that—while avoiding the pitfalls of significance testing—uses familiar statistical concepts: means, mean differences, and error bars. More importantly, it focuses on the effect size of one’s experiment/intervention, as opposed to significance testing (Claridge-Chang and Assam 2016). In comparison to typical NHST plots, estimation graphics have the following five significant advantages such as (1) avoid false dichotomy, (2) display all observed values (3) visualize estimate precision (4) show mean difference distribution. And most importantly (5) by focusing attention on an effect size, the difference diagram encourages quantitative reasoning about the system under study (Ho et al. 2019). Thus, we conducted a reanalysis of all of our two group data sets using both standard t tests and estimate statistics. In 2019, the Society for Neuroscience journal eNeuro instituted a policy recommending the use of estimation graphics as the preferred method for data presentation (Bernard 2021).

## ACKNOWLEDGEMENTS

We are very appreciative to the colleagues who supplied us with several fly strains. We thank Dr. Paul M. Macdonal (University of Texas at Austin) for sharing *UAS-bru2* line. We extend our gratitude to Drs. Yuh Nung and Lily Jan (UCSF) for their invaluable support and engagement in discussions pertaining to this project. This research was supported by a Startup funds from HIT Center for Life Science to WJK.

## DECLARATION OF GENERATIVE AI AND AI-ASSISTED TECHNOLOGIES IN THE WRITING PROCESS

During the creation of this work, the author(s) utilized QuillBot to rephrase English sentences, verify English grammar, and detect plagiarism, as none of the authors of this paper are native English speakers. After using this tool/service, the author(s) reviewed and edited the content as needed and take(s) full responsibility for the content of the publication.

## AUTHOR CONTRIBUTIONS

XZ and YH designed and performed revision experiments and revised manuscript. JB designed and performed experiments. WJK designed and performed experiments, wrote, and revised manuscript, analyzed data, and performed image analysis.

## REFERENCES

Aksoy-Aksel A, Zampa F, Schratt G. 2014. MicroRNAs and synaptic plasticity—a mutual relationship. Philos Trans R Soc B: Biol Sci. 369(1652):20130515. doi:10.1098/rstb.2013.0515.

Bartel DP. 2009. MicroRNAs: Target Recognition and Regulatory Functions. Cell. 136(2):215–233. doi:10.1016/j.cell.2009.01.002.

Bath E, Bowden S, Peters C, Reddy A, Tobias JA, Easton-Calabria E, Seddon N, Goodwin SF, Wigby S. 2017. Sperm and sex peptide stimulate aggression in female Drosophila. Nat Ecol Evol. 1(6):0154. doi:10.1038/s41559-017-0154.

Bernard C. 2021. Estimation Statistics, One Year Later. eNeuro. 8(2):ENEURO.0091-21.2021. doi:10.1523/eneuro.0091-21.2021.

Biryukova I, Asmar J, Abdesselem H, Heitzler P. 2009. Drosophila mir-9a regulates wing development via fine-tuning expression of the LIM only factor, dLMO. Dev Biol. 327(2):487–496. doi:10.1016/j.ydbio.2008.12.036.

Brodersen P, Voinnet O. 2009. Revisiting the principles of microRNA target recognition and mode of action. Nat Rev Mol Cell Biol. 10(2):141–148. doi:10.1038/nrm2619.

Cassidy JJ, Jha AR, Posadas DM, Giri R, Venken KJT, Ji J, Jiang H, Bellen HJ, White KP, Carthew RW. 2013. miR-9a Minimizes the Phenotypic Impact of Genomic Diversity by Buffering a Transcription Factor. Cell. 155(7):1556–1567. doi:10.1016/j.cell.2013.10.057.

Cassidy JJ, Straughan AJ, Carthew RW. 2015. Differential Masking of Natural Genetic Variation by miR-9a in Drosophila. Genetics. 202(2):675–687. doi:10.1534/genetics.115.183822.

Chapman T, Bangham J, Vinti G, Seifried B, Lung O, Wolfner MF, Smith HK, Partridge L. 2003. The sex peptide of Drosophila melanogaster: Female post-mating responses analyzed by using RNA interference. Proc National Acad Sci. 100(17):9923–9928. doi:10.1073/pnas.1631635100.

Claridge-Chang A, Assam PN. 2016. Estimation statistics should replace significance testing. Nat Methods. 13(2):108–109. doi:10.1038/nmeth.3729.

Cohen JE, Lee PR, Chen S, Li W, Fields RD. 2011. MicroRNA regulation of homeostatic synaptic plasticity. Proc Natl Acad Sci. 108(28):11650–11655. doi:10.1073/pnas.1017576108.

Corbel Q, Londoño Nieto C, Carazo P. 2022. Does perception of female cues modulate male short term fitness components in Drosophila melanogaster? Ecol Evol. 12(9):e9287. doi:10.1002/ece3.9287.

Daniel SG, Russ AD, Guthridge KM, Raina AI, Estes PS, Parsons LM, Richardson HE, Schroeder JA, Zarnescu DC. 2017. miR-9a mediates the role of Lethal giant larvae as an epithelial growth inhibitor in Drosophila. Biol Open. 7(1):bio027391. doi:10.1242/bio.027391.

Gallicchio L, Griffiths-Jones S, Ronshaugen M. 2020. Single-cell visualization of mir-9a and Senseless co-expression during Drosophila melanogaster embryonic and larval peripheral nervous system development. G3. 11(1):jkaa010. doi:10.1093/g3journal/jkaa010.

Gangadharan V, Kuner R. 2013. Pain hypersensitivity mechanisms at a glance. Dis Model Mech. 6(4):889–895. doi:10.1242/dmm.011502.

Gargano JW, Martin I, Bhandari P, Grotewiel MS. 2005. Rapid iterative negative geotaxis (RING): a new method for assessing age-related locomotor decline in Drosophila. Exp Gerontol. 40(5):386–95. doi:10.1016/j.exger.2005.02.005.

Gligorov D, Sitnik JL, Maeda RK, Wolfner MF, Karch F. 2013. A Novel Function for the Hox Gene Abd-B in the Male Accessory Gland Regulates the Long-Term Female Post-Mating Response in Drosophila. Plos Genet. 9(3):e1003395. doi:10.1371/journal.pgen.1003395.

Grishagin IV. 2015. Automatic cell counting with ImageJ. Anal Biochem. 473:63–65. doi:10.1016/j.ab.2014.12.007.

Gu P, Wang F, Shang Y, Liu J, Gong J, Xie W, Han J, Xiang Y. 2022. Nociception and hypersensitivity involve distinct neurons and molecular transducers in Drosophila. Proc Natl Acad Sci. 119(12):e2113645119. doi:10.1073/pnas.2113645119.

Häsemeyer M, Yapici N, Heberlein U, Dickson BJ. 2009. Sensory Neurons in the Drosophila Genital Tract Regulate Female Reproductive Behavior. Neuron. 61(4):511–518. doi:10.1016/j.neuron.2009.01.009.

Ho J, Tumkaya T, Aryal S, Choi H, Claridge-Chang A. 2019. Moving beyond P values: data analysis with estimation graphics. Nat Methods. 16(7):565–566. doi:10.1038/s41592-019-0470-3.

Hollis B, Koppik M, Wensing KU, Ruhmann H, Genzoni E, Erkosar B, Kawecki TJ, Fricke C, Keller L. 2019. Sexual conflict drives male manipulation of female postmating responses in Drosophila melanogaster. Proc Natl Acad Sci United States Am. 116(17):8437–8444. doi:10.1073/pnas.1821386116.

Hussain A, Üçpunar HK, Zhang M, Loschek LF, Kadow ICG. 2016. Neuropeptides Modulate Female Chemosensory Processing upon Mating in Drosophila. PLoS Biol. 14(5):e1002455. doi:10.1371/journal.pbio.1002455.

Isaacs D, Riordan H. 2020. Sensory hypersensitivity in Tourette syndrome: A review. Brain Dev. 42(9):627–638. doi:10.1016/j.braindev.2020.06.003.

Itai T, Hamanaka K, Sasaki K, Wagner M, Kotzaeridou U, Brösse I, Ries M, Kobayashi Y, Tohyama J, Kato M, et al. 2021. De novo variants in CELF2 that disrupt the nuclear localization signal cause developmental and epileptic encephalopathy. Hum Mutat. 42(1):66–76. doi:10.1002/humu.24130.

Katti P, Thimmaya D, Madan A, Nongthomba U. 2017. Overexpression of miRNA-9 Generates Muscle Hypercontraction Through Translational Repression of Troponin-T in Drosophila melanogaster Indirect Flight Muscles. G3: Genes, Genomes, Genet. 7(10):3521–3531. doi:10.1534/g3.117.300232.

Kozomara A, Birgaoanu M, Griffiths-Jones S. 2019. miRBase: from microRNA sequences to function. Nucleic Acids Res. 47(D1):D155–D162. doi:10.1093/nar/gky1141.

Kubli E. 2003. Sex-peptides: seminal peptides of the Drosophila male. Cell Mol Life Sci Cmls. 60(8):1689–1704. doi:10.1007/s00018-003-3052.

Latremoliere A, Woolf CJ. 2009. Central Sensitization: A Generator of Pain Hypersensitivity by Central Neural Plasticity. J Pain. 10(9):895–926. doi:10.1016/j.jpain.2009.06.012.

Lee SG, Sun D, Miao H, Wu Z, Kang C, Saad B, Nguyen K-NH, Guerra-Phalen A, Bui D, Abbas A-H, et al. 2023. Taste and pheromonal inputs govern the regulation of time investment for mating by sexual experience in male Drosophila melanogaster. PLOS Genet. 19(5):e1010753. doi:10.1371/journal.pgen.1010753.

Li Y, Wang F, Lee J-A, Gao F-B. 2006. MicroRNA-9a ensures the precise specification of sensory organ precursors in Drosophila. Gene Dev. 20(20):2793–2805. doi:10.1101/gad.1466306.

Li Z, Lu Y, Xu X-L, Gao F-B. 2013. The FTD/ALS-associated RNA-binding protein TDP-43 regulates the robustness of neuronal specification through microRNA-9a in Drosophila. Hum Mol Genet. 22(2):218–225. doi:10.1093/hmg/dds420.

Miao H, Wei Y, Lee SG, Wu Z, Kaur J, Kim WJ. 2024. Glia specific expression of neuropeptide receptor Lgr4 regulates development and adult physiology in Drosophila. J Neurosci Res. 102(1). doi:10.1002/jnr.25271.

Misra C, Bangru S, Lin F, Lam K, Koenig SN, Lubbers ER, Hedhli J, Murphy NP, Parker DJ, Dobrucki LW, et al. 2020. Aberrant Expression of a Non-muscle RBFOX2 Isoform Triggers Cardiac Conduction Defects in Myotonic Dystrophy. Dev Cell. 52(6):748–763.e6. doi:10.1016/j.devcel.2020.01.037.

Mohammadi AH, Seyedmoalemi S, Moghanlou M, Akhlagh SA, Zavareh SAT, Hamblin MR, Jafari A, Mirzaei H. 2022. MicroRNAs and Synaptic Plasticity: From Their Molecular Roles to Response to Therapy. Mol Neurobiol. 59(8):5084–5102. doi:10.1007/s12035-022-02907-2.

Öztürk-Çolak A, Marygold SJ, Antonazzo G, Attrill H, Goutte-Gattat D, Jenkins VK, Matthews BB, Millburn G, Santos G dos, Tabone CJ, et al. 2024. FlyBase: updates to the Drosophila genes and genomes database. GENETICS. 227(1):iyad211. doi:10.1093/genetics/iyad211.

Parrish JZ, Kim MD, Jan LY, Jan YN. 2006. Genome-wide analyses identify transcription factors required for proper morphogenesis of Drosophila sensory neuron dendrites. Genes Dev. 20(7):820–835. doi:10.1101/gad.1391006.

Petersen-Felix S, Curatolo M. 2002. Neuroplasticity - an important factor in acute and chronic pain. Swiss Méd Wkly. 132(2122):273–278. doi:10.4414/smw.2002.09913.

Pinho-Ribeiro FA, Verri WA, Chiu IM. 2017. Nociceptor Sensory Neuron–Immune Interactions in Pain and Inflammation. Trends Immunol. 38(1):5–19. doi:10.1016/j.it.2016.10.001.

Ram KR, Wolfner MF. 2007. Sustained Post-Mating Response in Drosophila melanogaster Requires Multiple Seminal Fluid Proteins. Plos Genet. 3(12):e238. doi:10.1371/journal.pgen.0030238.

Ren K, Dubner R. 2008. Neuron–glia crosstalk gets serious: role in pain hypersensitivity. Curr Opin Anaesthesiol. 21(5):570–579. doi:10.1097/aco.0b013e32830edbdf.

Salter MW. 2010. The Neurobiology of Central Sensitization. J Musculoskelet Pain. 10(1–2):23–33. doi:10.1300/j094v10n01_03.

Smalheiser NR, Lugli G. 2009. microRNA Regulation of Synaptic Plasticity. NeuroMolecular Med. 11(3):133–140. doi:10.1007/s12017-009-8065-2.

Subramanian M, Hyeon SJ, Das T, Suh YS, Kim YK, Lee J-S, Song EJ, Ryu H, Yu K. 2021. UBE4B, a microRNA-9 target gene, promotes autophagy-mediated Tau degradation. Nat Commun. 12(1):3291. doi:10.1038/s41467-021-23597-9.

Suh YS, Bhat S, Hong S-H, Shin M, Bahk S, Cho KS, Kim S-W, Lee K-S, Kim Y-J, Jones WD, et al. 2015. Genome-wide microRNA screening reveals that the evolutionary conserved miR-9a regulates body growth by targeting sNPFR1/NPYR. Nat Commun. 6(1):7693. doi:10.1038/ncomms8693.

Wang Y, Wang H, Li X, Li Y. 2016. Epithelial microRNA 9a regulates dendrite growth through Fmi Gq signaling in Drosophila sensory neurons. Dev Neurobiol. 76(2):225–237. doi:10.1002/dneu.22309.

Woolf CJ, Salter MW. 2000. Neuronal Plasticity: Increasing the Gain in Pain. Science. 288(5472):1765–1768. doi:10.1126/science.288.5472.1765.

Yang C, Rumpf S, Xiang Y, Gordon MD, Song W, Jan LY, Jan Y-N. 2009. Control of the Postmating Behavioral Switch in Drosophila Females by Internal Sensory Neurons. Neuron. 61(4):519–526. doi:10.1016/j.neuron.2008.12.021.

Yapici N, Kim Y-J, Ribeiro C, Dickson BJ. 2008. A receptor that mediates the post-mating switch in Drosophila reproductive behaviour. Nature. 451(7174):33–37. doi:10.1038/nature06483.

Yasunaga K, Tezuka A, Ishikawa N, Dairyo Y, Togashi K, Koizumi H, Emoto K. 2015. Adult Drosophila sensory neurons specify dendritic territories independently of dendritic contacts through the Wnt5–Drl signaling pathway. Genes Dev. 29(16):1763–1775. doi:10.1101/gad.262592.115.

Yatsenko AS, Shcherbata HR. 2014. Drosophila miR-9a Targets the ECM Receptor Dystroglycan to Canalize Myotendinous Junction Formation. Dev Cell. 28(3):335–348. doi:10.1016/j.devcel.2014.01.004.

Ye Y, Xu H, Su X, He X. 2016. Role of MicroRNA in Governing Synaptic Plasticity. Neural Plast. 2016:4959523. doi:10.1155/2016/4959523.

Yuva-Aydemir Y, Simkin A, Gascon E, Gao F-B. 2011. MicroRNA-9. RNA Biol. 8(4):557–564. doi:10.4161/rna.8.4.16019.

Zhang X, Sun D, Wong K, Salkini A, Najafi H, Kim WJ. 2024. The astrocyte-enriched gene deathstar plays a crucial role in the development, locomotion, and lifespan of D. melanogaster. Fly. 18(1):2368336. doi:10.1080/19336934.2024.2368336.

Zhu EY, Guntur AR, He R, Stern U, Yang C-H. 2014. Egg-Laying Demand Induces Aversion of UV Light in Drosophila Females. Curr Biol. 24(23):2797–2804. doi:10.1016/j.cub.2014.09.076.

