## Supplementary figures and images for "Hypersensitivity controlled by mir-9a modulates female receptivity of *Drosophila melanogaster*"

### Supplemental figure 1

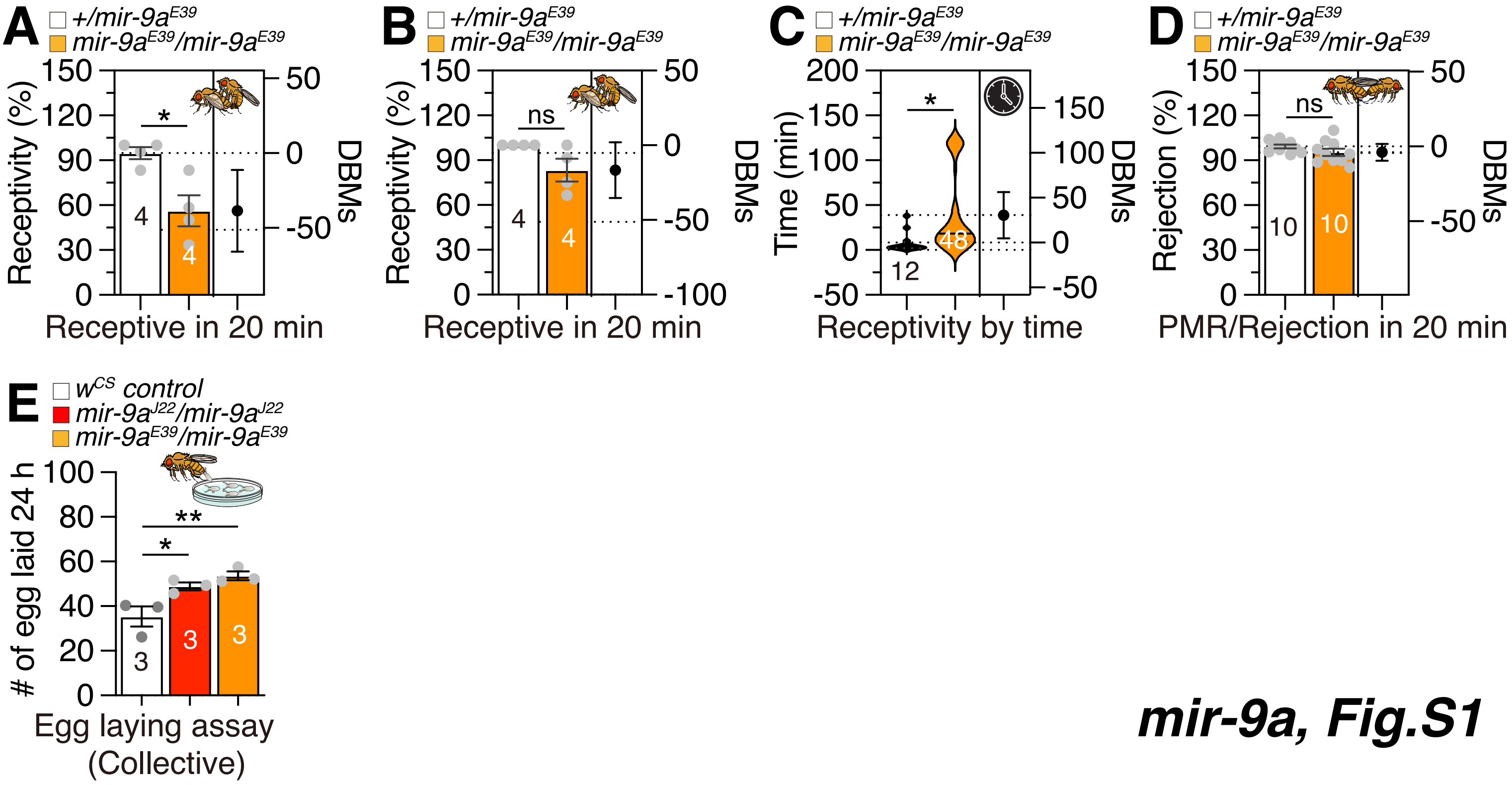

### Supplemental figure 2

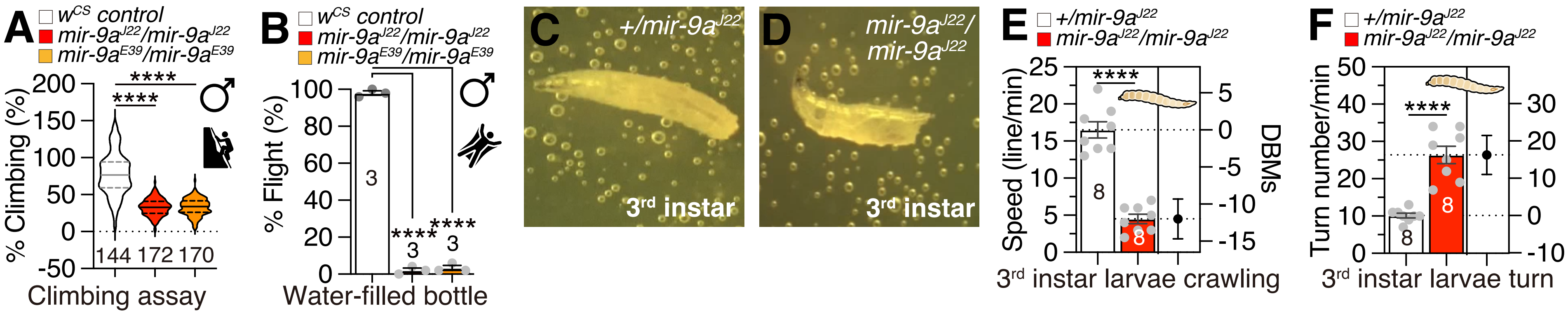

### Supplemental figure 3

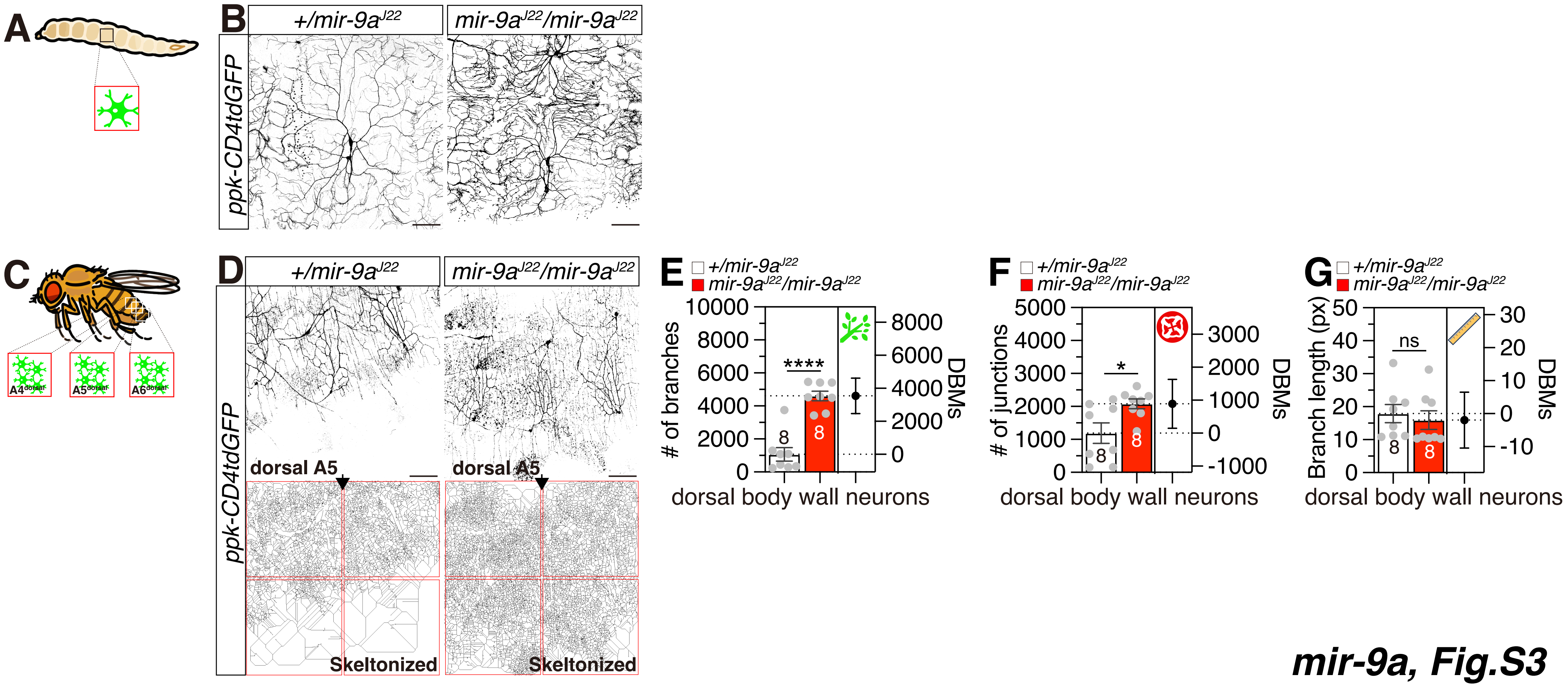

### Supplemental figure 4

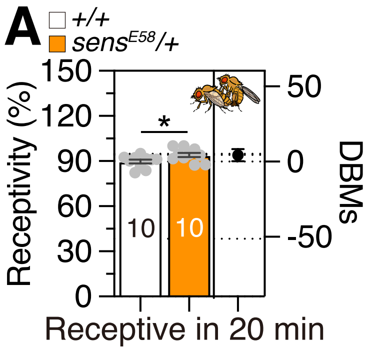
